# Oscillatory Mechanisms of Successful Memory Formation in Younger and Older Adults Are Related to Structural Integrity

**DOI:** 10.1101/530121

**Authors:** Myriam C. Sander, Yana Fandakova, Thomas H. Grandy, Yee Lee Shing, Markus Werkle-Bergner

## Abstract

We studied oscillatory mechanisms of memory formation in 48 younger and 51 older adults in an intentional associative memory task with cued recall. While older adults showed lower memory performance than young adults, we found subsequent memory effects (SME) in alpha/beta and theta frequency bands in both age groups. Using logistic mixed effect models, we investigated whether interindividual differences in structural integrity of key memory regions could account for interindividual differences in the strength of the SME. Structural integrity of inferior frontal gyrus (IFG) and hippocampus was reduced in older adults. SME in the alpha/beta band were modulated by the cortical thickness of IFG, in line with its hypothesized role for deep semantic elaboration. Importantly, this structure–function relationship did not differ by age group. However, older adults were more frequently represented among the participants with low cortical thickness and consequently weaker SME in the alpha band. Thus, our results suggest that differences in the structural integrity of the IFG contribute not only to interindividual, but also to age differences in memory formation.

---

Episodic memory, the ability to remember episodes with their spatial and temporal details and context (Tulving 2002) declines with age (Shing et al. 2010). Longitudinal studies have shown that the decline in memory performance starts on average after the age of 60, but with massive interindividual differences (Nyberg 2017). The causes of episodic memory decline are multifaceted (for a review, see Nyberg et al. 2012; Wang and Cabeza 2017), including structural and functional age-related changes in key memory regions, e.g., medio-temporal (MTL), prefrontal cortical (PFC) and parietal regions (Cabeza et al. 2008; Raz et al. 2005), white matter decline (Davis et al. 2009), changes in functional and structural connectivity (Fjell et al. 2016; Grady 2017; Davis et al. 2019), and declines in neurotransmitter systems (Bäckman et al. 2006; Mather and Harley, 2016). These senescent changes are hypothesized to underlie typically observed age differences in all stages of memory processing, for example, during encoding (Craik and Rose 2012), particularly with regard to the binding of associative information (Naveh-Benjamin, 2000), during retrieval (Wang and Cabeza, 2017; Fandakova et al. 2018), sleep-dependent consolidation (Helfrich et al. 2018; Muehlroth et al. 2019), and forgetting (Fandakova et al. 2019). Age differences in encoding may be particularly crucial, since differences in the quality of memory representations may have downstream consequences for later stages of memory processing such as consolidation and retrieval (see Fandakova et al. 2018). To date, it is still an open question how age differences in the structural integrity of memory-relevant brain regions relate to differences in their functional recruitment during memory encoding. Is memory-relevant activation modulated by structural integrity of key memory regions?

Mechanisms of successful memory formation can be studied with the *subsequent memory paradigm* (Paller and Wagner 2002; Werkle-Bergner et al. 2006). This approach makes use of the fact that not all encoded information can later be remembered. Comparing the neural dynamics in trials with stimuli that will subsequently be remembered against those that will subsequently not be remembered reveals the neural underpinnings of successful memory formation. Functional magnetic resonance imaging (fMRI) studies using this paradigm have provided convincing evidence for the contribution and interaction of MTL and PFC regions to successful memory formation (e.g., Wagner et al. 1998; Reber et al. 2002) as part of a broader episodic memory network (Benoit and Schacter 2015; Renoult et al. 2019). In particular, the MTL, and more specifically the hippocampus (HC), is regarded as crucial for binding pieces of information into a coherent memory representation, whereas PFC regions serve the selection and elaboration of encoded information (Miller and Cohen 2001; Simons and Spiers 2003). Within the PFC, prominent roles have been attributed to the left inferior frontal gyrus (IFG) for memory formation of semantic information and to the right IFG for memory formation of pictorial information (Paller and Wagner 2002).

Thus, episodic memory formation crucially depends on interactions between regions of the PFC and regions of the MTL (Simons and Spiers 2003). A mechanism for efficient representation and communication in broad neural networks is rhythmic neural activity (von der Malsburg 1995; Fries 2005; Parish et al. 2018). In particular, increases in oscillatory theta power and decreases in alpha/beta power support successful encoding of episodes (Hanslmayr and Staudigl 2014; but note that some studies have also reported theta decreases). On a cognitive level, alpha/beta oscillations seem to reflect elaborative encoding processes (Hanslmayr et al. 2012), whereas theta oscillations may serve associative binding of information (Clouter et al. 2017). This picture is completed by a study that simultaneously assessed SME in electroencephalography (EEG) and fMRI, and identified the IFG as the source region of SME in the alpha/beta band, and the MTL (together with the lateral temporal cortex) as the source region of SME in the theta band (Hanslmayr et al. 2011). A recent study using transcranial magnetic stimulation (TMS) of the left IFG even demonstrated a causal role of (beta) desynchronization for memory formation (Hanslmayr et al. 2014; see Rossi et al. 2011, for related evidence).

While several studies have compared SME in younger and older adults using fMRI (for a meta-analysis of 18 studies, see Maillet and Rajah 2014), surprisingly little is known about age differences in oscillatory neural mechanisms of episodic memory formation (Werkle-Bergner et al. 2006, but see Strunk and Duarte 2019). It seems reasonable to hypothesize that oscillatory neural activity observed with EEG parallels effects observed in BOLD activation levels of IFG and HC; however, this relation is not sufficiently established so far. To date, only one study investigated single-trial correlations between BOLD activation in the IFG and alpha SME and MTL activation and theta SME (Hanslmayr et al. 2011). The evidence from fMRI studies makes it difficult to predict whether one would observe differences in SME between younger and older adults when using oscillatory scalp-recorded EEG measures. In particular, BOLD level measures are only an indirect measure of neural activity and may be prone to confounds due to age differences in cardiovascular couplings (e.g., Gazzaley and D’Esposito 2005; Rugg, 2016). Here, we therefore examined to what extent patterns of oscillatory neural activity related to memory formation are altered in older adults as compared to younger adults, with regard to SME in the theta and alpha band. Furthermore, to the best of our knowledge, the relation between structural integrity of key regions of memory functions and oscillatory mechanisms of memory formation has not been investigated. While previous studies have shown <u>functional</u> activation of the IFG and HC during memory formation (mostly using fMRI, see Paller and Wagner 2002), studies have not directly related functional activation of the IFG or HC during memory encoding to their <u>structural</u> integrity. Second, other studies have provided evidence that the structural integrity of PFC and MTL is important for memory performance (e.g. HC gray matter being related to episodic memory performance, Gorbach et al. 2017; but see Van Petten, 2004), but have not shown that memory-relevant functional activation was modulated by the structural integrity of that region. Thus, so far, only bilateral relations between (a) functional activation of HC and IFG and memory performance and (b) structural integrity and memory performance have been established. Studies have also not yet closed the triadic loop linking measures of the structural integrity, functional activation, and memory performance to each other. We hypothesized that while theta and alpha power modulations (i.e., SME) explain accuracy on a trial-by-trial level, differences in MTL and PFC structure may be related to between-person differences in accuracy. As SME in the alpha band are thought to reflect elaborative processing of information, we hypothesized that SME in that range depend on the structural integrity of the IFG, which has previously been *functionally* related to subsequent memory (Hanslmayr et al. 2011). Second, as SME in the theta band are thought to reflect interactions between the HC and PFC (Klimesch, 1999; Nyhus and Curran 2010) and have been localized to the MTL and adjacent regions (Hanslmayr et al. 2011), we hypothesized that the degree of theta power modulation depends on the structural integrity of the MTL, in particular the HC. Importantly, since both MTL and PFC show pronounced structural and functional decline in normal aging (West, 1996; Raz et al. 2005; Shing et al. 2010), we expected large between-person differences in structural integrity in an age-comparative setting to be particularly conducive to delineate these structure–function relationships. We hypothesized that reduced structural integrity of HC and PFC in older adults would be accompanied by smaller SME in theta and alpha frequency bands in this age group.

We used repeated cued-recall tests with feedback to track learning of a large set of scene–word pairs in younger and older adults. Specifically, younger and older adults were instructed to study and try to remember scene–word pairs by forming an integrated mental image of the pair (cf. Fandakova et al. 2018; Muehlroth et al. 2019). Prior to study, all participants were instructed in an imagery strategy that has been shown to increase associative memory in younger and older adults effectively (Brehmer et al. 2007; Shing et al. 2008). In contrast to most other studies examining SME in older adults (for review, see Maillet and Rajah 2014) we used cued verbal recall instead of a recognition procedure to test memory. While correct responses in a cued-recall task depend on remembering the specific scene–word binding, performance on recognition tasks may also, at least partially, be supported by additional processes such as the overall familiarity of the presented scenes and words (Yonelinas, 2002). Older adults often learn more slowly (i.e., they need more repetitions to learn the same amount of information; e.g., Li et al. 2004) and they reach their limit of acquisition earlier than younger adults do (i.e., they remember fewer items; e.g., Rugg and Morcom 2005). Since SME analyses require both age groups to perform in a similar range, we opted for a task design that would eliminate or at least reduce age differences in memory performance, and would allow us to track the learning history of individual items within each participant (see Fig. 1). We therefore used different numbers of trials and different numbers of learning and recall cycles for younger and older adults (see Method for the details). We simultaneously recorded EEG while participants encoded and recalled scene–word pairs. In addition, we used structural MRI (sMRI) to assess IFG cortical thickness and HC volume. We examined the role of within-person power modulations and between-person differences in structure for the prediction of single trial accuracy, modelling them simultaneously in a logistic mixed effects model. We hypothesized that oscillatory mechanisms of memory formation depend on the structural integrity of HC and IFG, which are affected by advancing age, thus leading to less successful encoding in older compared to younger adults.

**Figure 1.**
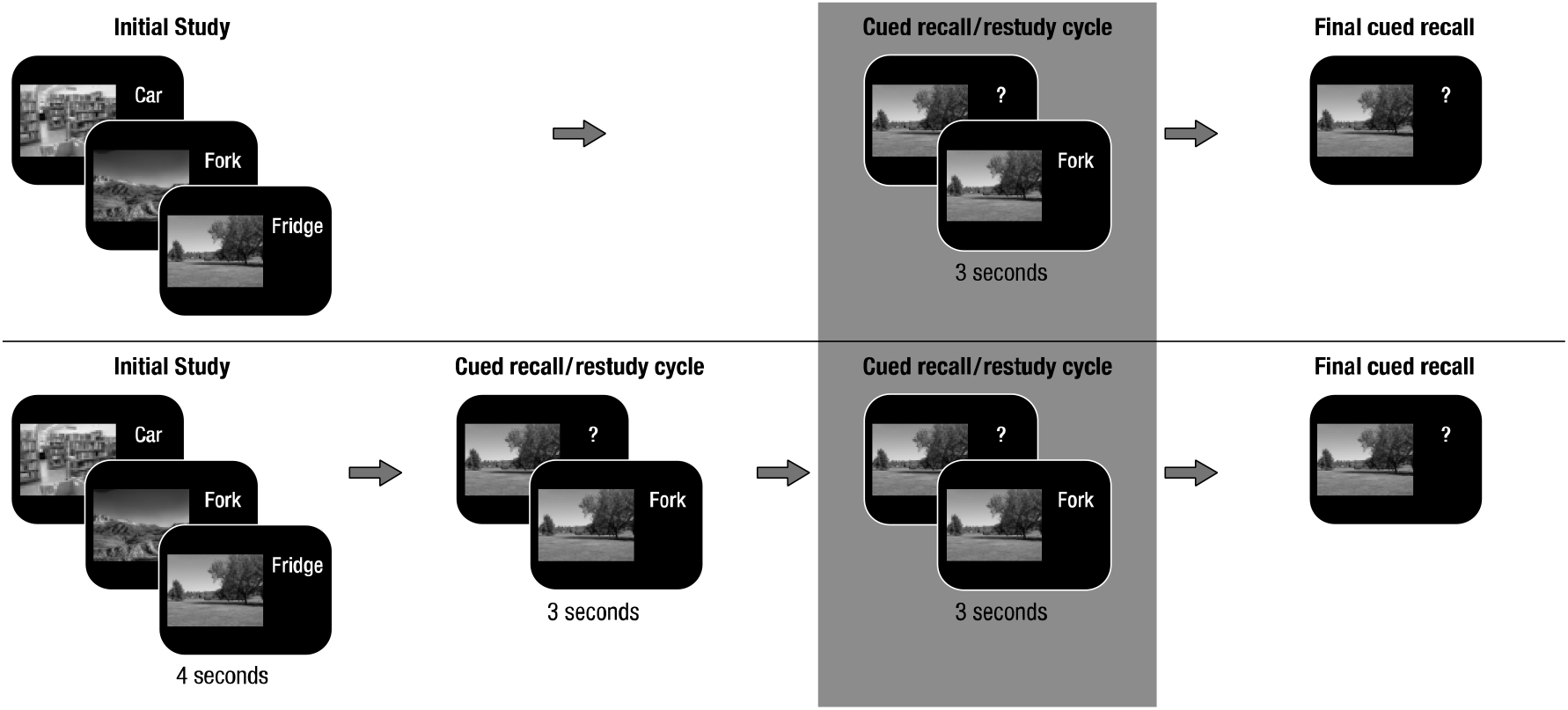
Memory Paradigm (cf. Fandakova et al. 2018; Muehlroth et al. 2019). (A) During initial study, participants were instructed to remember 440 scene–word pairs (younger adults) or 280 scene–word pairs (older adults). (B) During the cued recall/restudy phase, the scene was presented as a cue to recall the corresponding word. Irrespective of recall accuracy, the original pair was presented again to allow for restudy. The whole cued recall/restudy cycle was performed once in younger adults and twice in older adults. (C) During final recall, scenes again served as cues to recall the corresponding word, but no occasion for restudy was provided. Subsequent memory analysis was done on the last cued restudy of the scene–word pairs before final cued recall (marked by the grey background).

## Materials and Methods

The present data were derived from a series of studies investigating age-related differences in the encoding, consolidation, and retrieval of associative memories (see Fandakova et al. 2018, for the effects of age and memory quality on false memory retrieval and Muehlroth et al. 2019, for the effects of sleep on memory retrieval). At the core of the experimental design was an associative scene–word pair memory paradigm, consisting of a learning session on the first day (Day 1) and a delayed recognition or delayed cued-recall task approximately 24 hours later (Day 2) (see Figure 1 for a depiction of the study procedure of Day 1). We collected sMRI and fMRI data during and after delayed recall or recognition on Day 2. In part of the sample, sleep was also monitored at participants’ homes using ambulatory polysomnography (PSG). As the current study focusses on age differences in encoding (Day 1), neither fMRI nor PSG data are included in the present report (see Fandakova et al. 2018; Muehlroth et al. 2019, respectively). We included sMRI data to test our hypothesis that structural integrity of the HC (i.e., HC volume) and IFG (i.e., cortical thickness) may be related to oscillatory SME in theta and alpha frequencies.

### Participants

Data from participants in the two studies (cf. Fandakova et al. 2018; Muehlroth et al. 2019) were jointly processed in the present analyses. While the procedure of these two studies differed with regard to subsequent tests on the following days, the learning procedure on Day 1 was identical in both, except for the pairing of scene–word combinations (see below for more information). Hence, the data were combined and analyzed here. In total, data from 141 participants (61 younger and 80 older adults) were available. All participants were right-handed native German speakers, had normal or corrected-to-normal vision, no history of psychiatric or neurological disease, and did not take psychiatric medication. Younger adults were students enrolled in local universities. Due to technical failures or extreme artifacts, EEG data were only available for 114 participants (50 younger and 63 older adults). This sample was used for the determination of time-frequency clusters with SME on the grand-average EEG data.

In the EEG sample, 3 younger and 11 older adults did not provide full MRI data sets (T1 and/or T2 missing or containing strong motion artefacts). The effective final sample in the main analyses comprised 48 younger adults (M_age_ (SD) = 24.19 (2.54), range 19.12–27.87 years, 24 females), and 51 older adults (M_age_ (SD) = 70.18 (2.75), range 63.78–75.75 years, 28 females). Performance of the final sample did not differ from the EEG-only sample. Older adults were screened with the Mini-Mental State Exam (MMSE; Folstein et al. 1975) and none had a value below the threshold of 26 points. All participants were assessed on marker tests of verbal knowledge (Spot-a-Word, cf. Lehrl, 1977) and perceptual speed (digit symbol substitution test, cf. Wechsler, 1955) and showed age-typical performance (younger adults: M_Digit Symbol_ (SD) = 69.15 (10.66), M_Spot-a-word_(SD) = 23.40 (3.18), older adults: M_Digit Symbol_ (SD) = 50.84 (10.71), M_Spot-a-word_(SD) = 29.10 (3.19), with younger adults showing higher levels of perceptual speed, *t*(97) =8.51, *p* < .001, than older adults, and older adults showing higher levels of verbal knowledge, *W* = 232, *p* < .001, than younger adults. Given that the two study designs were both physically demanding and time-consuming (EEG and MR sessions across several days), the older adult sample in particular is certainly not representative for their age group, but rather represents individuals aging healthily. However, performance in the marker tasks was similar to other cognitive neuroscience studies previously run in our research center (e.g., Sander et al. 2011; Fandakova et al. 2014) and showed a typical pattern of age differences. In order to ensure comparability of younger and older adults in our sample, we also only selected university students, i.e., younger adults who are most likely to become highly educated and healthy older adults. Information on 45 young adults’ and 49 older adults’ educational level was available. The highest school degree was the “Abitur” (highest school certificate in Germany that allows entrance to university education), which was obtained by 32 young adults and 9 older adults, and 13 of the younger and 24 of the older adults had obtained a university degree. Young adults had spent more years in school, *M(SD)* = 12.71(0.73), than older adults, *M(SD)* = 11.70(1.58), *W* = 1589, *p* < .001, but older adults had spent more time at university (YA: *M(SD)* = 3.15 (2.59), OA: *M(SD)* = 5.17 (2.35), *W* = 446.5 *p* < .001). No matching between age groups was performed on any of the demographic variables. The ethics committee of the Deutsche Gesellschaft für Psychologie (DGP) approved the study.

### Experimental Paradigm

A subsequent memory paradigm (Paller and Wagner 2002) was used to compare neural oscillations related to later remembered versus later not-remembered items. Initially, participants were instructed to memorize randomly paired scene–word stimuli using an imagery strategy. Participants were strongly encouraged to generate integrated mental images of the pairs that were vivid and creative. Examples were discussed in detail until the strategy was well understood. Previous studies have provided evidence that without instructions, younger and older adults spontaneously adopt different encoding strategies. Young adults often elaborate on items, whereas older adults employ rote-based strategies (see Rugg and Morcom 2005, for this line of argumentation). Differences in encoding strategies (e.g., shallow versus deep encoding) have been shown to modulate SME (Hanslmayr and Staudigl 2014) and age-related neural differences are often confounded with differences in cognitive operations or strategies engaged by younger and older adults. The explicit strategy instruction in our study therefore aimed to minimize this confound and gain control over participants’ learning strategy.

During the experiment, scene–word pairs were presented for 4 seconds, with the scene on the left and the word on the right of the screen. During this initial presentation, participants used a four-point imaginability scale to indicate how well they were able to form an integrated image of the scene and word. In subsequent blocks, the scenes served as cues and participants had to verbally recall the associated word. Verbal responses were digitally recorded. Recall time was not constrained. The accuracy of the answers was coded online by the experimenter. Independent of recall accuracy, the correct word was shown again together with the scene (for 3 seconds), fostering further learning of the pair. Then participants completed a final cued-recall task without feedback. As differences in performance level have been shown to confound age differences in neural activity (Rugg 2017), we adjusted task difficulty between the two age groups in the following way. (1) Since older adults’ memory performance was expected to be generally lower, younger adults learned 440 pairs, whereas older adults learned 280 pairs. (2) Older adults completed an additional cued-recall/restudy cycle before the final test, in line with an approach implemented by other studies that used different numbers of encoding cycles in younger and older adults (Daselaar et al. 2006; Morcom et al. 2007; Duverne et al. 2008). With this age-adapted study design, we aimed to identify age-related differences in brain activity associated with memory formation that are free of the influence of confounding variables such as performance differences between age groups (Rugg and Morcom 2005). While younger adults were able to learn a number of scene–word pairs that would allow for subsequent memory analysis of the initial study phase, older adults’ initial performance was too low for such an analysis (but see Sommer et al. 2019, for an alternative age-comparative analysis of this initial study phase). Therefore, the subsequent memory analysis of the EEG data in both age groups is focused on the last restudy phase before the final test. Pairs recalled correctly prior to this last encoding phase were omitted from the analysis.

During the experimental procedure, participants were seated comfortably in a dimly lit room that was electromagnetically and acoustically shielded. The EEG measurement started with a 6-minute relaxation phase (resting EEG), followed by the task. Between blocks, participants were allowed to take breaks and leave the cabin.

### Stimuli

Stimuli are described in detail by Fandakova et al. (2018). Briefly, we selected 580 picture stimuli, half of them depicting indoor scenes and the other half depicting outdoor scenes. In addition, 580 concrete nouns with 2 phonetic syllables and a word length of 4–8 letters were selected from the CELEX database of the Max Planck Institute for Psycholinguistics (http://celex.mpi.nl/). Pictures and words were randomly paired to form stimuli for the presentation during the experiment. Note that there was a difference between the two original studies in how stimuli were paired: Whereas all participants of the first sample saw the same pairings, each participant of the second sample saw a different combination of pairings.

### Analysis of Behavioral Data

Behavioral data was analysed using R 3.5.2 (R Development Core Team, 2018). Raincloud plots were used for illustration of the data (Allen et al. 2019).

#### Performance Across the Whole Learning Procedure

Performance for each recall cycle (i.e., 2 for younger adults, and 3 for older adults) was calculated as the proportion of correctly recalled items out of all presented items (i.e., 440 for younger adults and 280 for older adults).

#### Overall Learning Success

Overall learning success refers to the proportion of correctly recalled items out of all presented items at the last final cued-recall task and was compared between age groups using the Wilcoxon rank sum test since assumptions of normality were violated.

#### Learning Gain in the Last Restudy Phase

To keep the behavioral analysis in line with the subsequent memory analysis, our main behavioral measure of interest was the *learning gain in the last restudy phase*. We therefore computed the learning gain as the percentage of items correctly recalled in the final cued recall out of those pairs that were not previously recalled in earlier recall cycles. Differences in learning gains were compared between age groups using the Wilcoxon rank sum test.

#### Imagery Ratings

Participants rated the imaginability of each scene–word pair during the initial study phase. Unfortunately, these imagery ratings contained many missing trials, mostly because of a technical programming mistake. Post-hoc inspection of our data revealed that participants often seem to have run out of time for the imagery rating and to have given their rating too late for registration or even during the next trial. This led to missing data in the following trial since only one response was registered per trial. We therefore excluded trials with missing responses or reaction times below 500 ms from the analysis. The number of trials included in the analysis was *M (SD)* = 222.58 (75.37) in the younger and *M (SD)* = 134.02 (59.99) in the older adults.

To investigate whether adults of both age groups were able to modulate their imagery ratings according to subsequent memory success, we compared imagery ratings for later recalled and not-remembered trials within each age group using Wilcoxon signed-rank tests. We then tested whether age groups differed in the modulation of the imagery ratings by comparing individual difference values (remembered minus not-remembered pairs) between age groups using the Wilcoxon rank sum test.

### Acquisition and structural MR analyses

Whole-brain MRI data were acquired on a Siemens Magnetom 3T Tim Trio scanner. A high-resolution T1-weighted MPRAGE sequence (TR = 2500 ms, TE = 4.77 ms, FOV = 256 mm, voxel size = 1 × 1 × 1 mm^3^) was collected from each participant. Cortical thickness was estimated using Freesurfer 5.1.0 following the Freesurfer standard image analysis processing pipeline as described on (http://surfer.nmr.mgh.harvard.edu/). This pipeline generates assessments of cortical thickness, calculated as the closest distance from the gray/white boundary to the gray/CSF boundary at each vertex on the tessellated surface (Fischl and Dale, 2000). Parcellation of the cerebral cortex into units with respect to gyral and sulcal structure was performed using the Desikan-Atlas (Desikan et al. 2006). Cortical thickness per subject was extracted for pars triangularis, pars orbitalis, and pars opercularis separately for the left and the right hemisphere. Following previous studies examining age differences in cortical thickness in aging (e.g., Fjell et al. 2009 for a large multi-site study), a trained observer manually checked the accuracy of the spatial registration and the white matter and gray matter segmentations. One participant was excluded due to registration problems. To capture the structural integrity of the IFG for a given person, we computed the sum of cortical thickness of these six regions (i.e., collapsing across hemispheres).

Since the automatic procedure pipeline in Freesurfer has been shown to selectively overestimate hippocampal volume in younger adults and thereby to bias age comparisons (Wenger et al. 2014), we acquired images of the MTL using a high-resolution, T2-weighted 2D turbo-spin echo (TSE) sequence, oriented perpendicularly to the long axis of the hippocampus (in-plane resolution: 0.4 mm × 0.4 mm, slice thickness: 2 mm, 31 slices, image matrix: 384 × 384, TR: 8150 ms, TE: 50 ms, flip angle: 120°) that was optimized for hippocampal subfield volume estimation (cf. Keresztes et al. 2017; Shing et al. 2011). Total volume of the hippocampal body was estimated as the sum of HC subfields including CA1, dentate gyrus, and subiculum and corrected for intracranial volume. The subfields were segmented using a semi-automated procedure with a custom-built hippocampal subfield atlas (both the procedure and the atlas are described in Bender et al. (2018) applying ASHS (Automatic Segmentation of Hippocampal Subfields; Yushkevich et al. 2015). Since the functional connection between the hippocampal subfields are strong, but poorly understood, and the specific contribution of single subfields to scalp-measured theta SME has not yet been established, we took the sum of left and right HC total volume of all subfields, thus the total HC body, as a measure of HC structural integrity. Differences in structural integrity were compared between age groups using independent sample t-tests.

### EEG Recording and Preprocessing

EEG was recorded continuously with BrainVision amplifiers (BrainVision Products GmbH, Gilching, Germany) from 61 Ag/Ag-Cl electrodes embedded in an elastic cap. Three additional electrodes were placed at the outer canthi and below the left eye to monitor eye movements. During recording, all electrodes were referenced to the right mastoid electrode, while the left mastoid electrode was recorded as an additional channel. The EEG was recorded with a band-pass of 0.1 to 250 Hz and digitized with a sampling rate of 1000 Hz. During preparation, electrode impedances were kept below 5 kΩ.

EEG data preprocessing was performed with the Fieldtrip software package (developed at the F. C. Donders Centre for Cognitive Neuroimaging, Nijmegen, The Netherlands; http://fieldtrip.fcdonders.nl/) supplemented by custom-made MATLAB code (The MathWorks Inc., Natick, MA, USA). An independent-component analysis (ICA) was used to correct for eye blink and cardio artifacts (Jung et al. 2000). Independent components representing artifactual sources were automatically detected, visually checked, and removed from the data. For analyses, the EEG was demeaned, re-referenced to mathematically linked mastoids, band-pass filtered (0.2–100 Hz; fourth order Butterworth) and downsampled to 250 Hz. Automatic artifact correction was performed for remaining artifacts following the FASTER procedure (Nolan et al. 2010). Excluded channels were interpolated with spherical splines (Perrin et al. 1989). In order to facilitate comparison with other studies, data was re-referenced to an average reference after artifact correction.

Data epochs were selected from the last cued-recall/restudy cycle. Four-second data epochs were extracted from −1 s to 3 s with respect to the onset of scene–word presentation during the last restudy phase. Time-frequency representations (TFRs) within the frequency range of interest (2–20 Hz) were derived from a short-time Fourier analysis with Hanning tapers with a fixed width of 500 ms, resulting in frequency steps of 2 Hz. Single-trial power was log-transformed. Only trials with *subsequently* remembered or not-remembered stimuli were included in the analysis.

In order to account for differences between age groups in procedure prior to the last restudy phase, trials with stimuli that were successfully remembered prior to the final cued recall/restudy cycle were omitted from the analysis. The number of trials included was *M(SD)* = 290.50 (42.55) for younger adults and *M(SD)* = 183.33 (39.52) for older adults.

### Analysis of Oscillatory Activity at the Group Level

First, trials with stimuli that were remembered and trials with stimuli that were not remembered during the final cued recall were averaged for each subject. We then determined time-frequency clusters on the grand-average level (collapsed across age groups) that showed reliable differences between subsequently remembered and not-remembered trials. Note that each participant is entered into this analysis with average data that are not weighted for the number of trials and therefore do not bias the analysis towards participants with more or fewer trials. More specifically, as the clusters were determined independently of age group, differences in trial numbers between the age groups did not affect this analysis. In subsequent analyses of the role of structural integrity of IFG and HC for single-trial power modulations within the identified time-frequency clusters, we use a mixed effect model with a random subject factor which accounts for between-person differences in trial numbers. We used dependent-sample t-tests on all electrodes across the whole trial length (from stimulus onset to 3 s). The threshold for electrodes to be included in a cluster was set to *p* = .05 and clusters were defined as a minimum of two neighboring electrodes showing reliable differences in activity. We controlled for multiple comparisons using non-parametric cluster-based permutation tests (Maris and Oostenveld 2007). The permutation null-distribution for the resulting t-values was determined by randomly switching the condition labels 1000 times and recomputing the t-tests. Note that we excluded one younger and two older adults’ data with final recall accuracy below 10% or above 90% from this part of the analysis in order not to bias the results via participants with highly unbalanced trial numbers across conditions. However, these participants were only excluded for the determination of the clusters of interest, but included in the subsequent data modelling as our mixed effects model with a random subject factor was able to account for differences in trial numbers.

To explain the next steps, we need to foreshadow the results of this analysis: The cluster-based permutation statistics yielded two significant (*p* < .025) clusters of electrodes that were considered as regions of interest in subsequent analyses (see Figure 2). The early cluster had a maximum around 500–800 ms and was predominantly found in the theta frequency range (4–6 Hz). The later cluster had a maximum around 1000–2000 ms and encompassed alpha and beta frequencies (8–20 Hz). To ease comprehension, we will refer to the earlier cluster as the SME in the theta band and to the later cluster as the SME in the alpha/beta frequency band.

**Figure 2.**
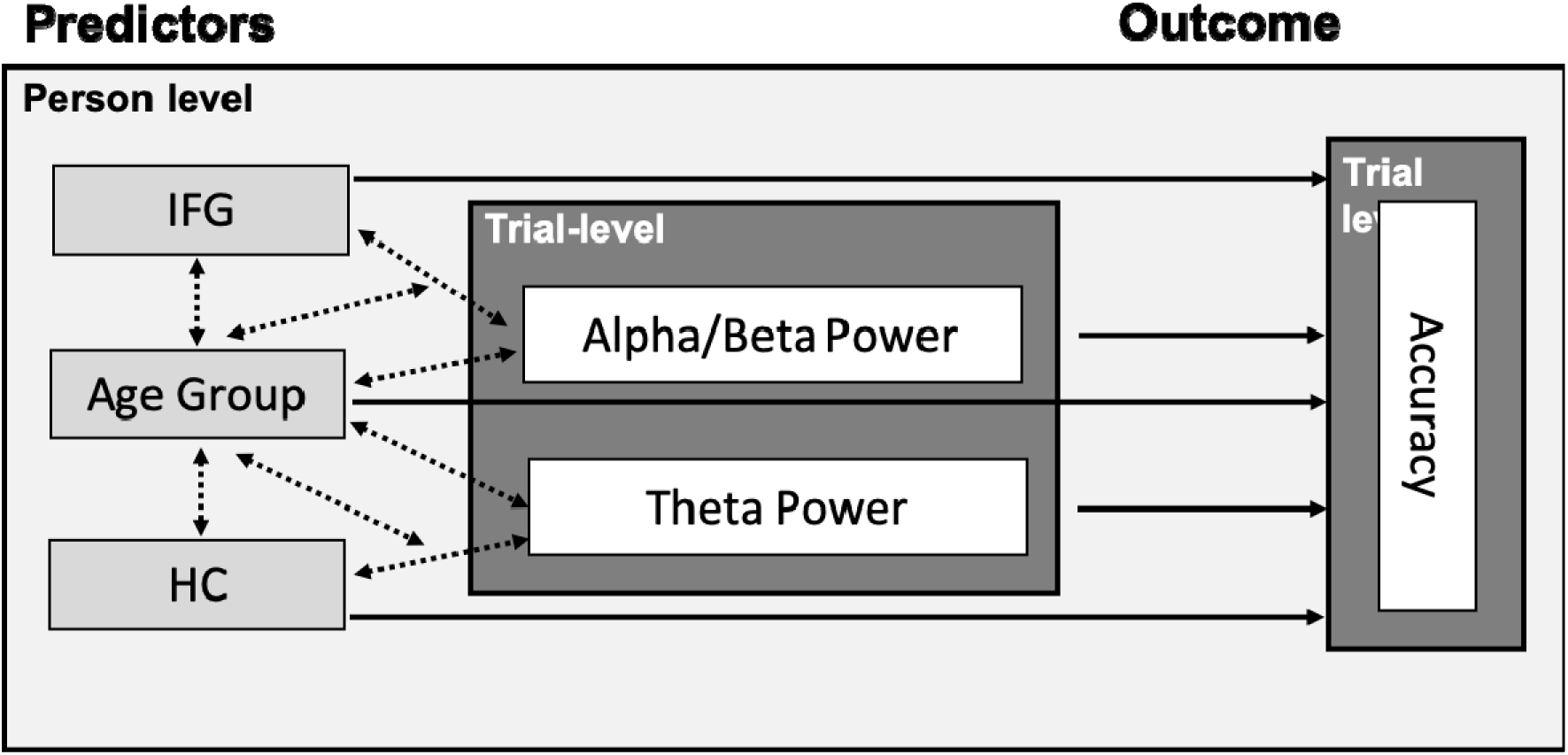
IIIustration of the mixed-effects model. On the trial level, alpha/beta power and theta power are used to predict single-trial accuracy. Person-level predictors are measures of structural integrity (IFG and HC) and age group. Tested main effects are represented by solid lines and single-headed arrows, and interactions by dotted lines and double-headed arrows. IFG: inferior frontal gyrus; HC: hippocampus.

**Figure 3.**
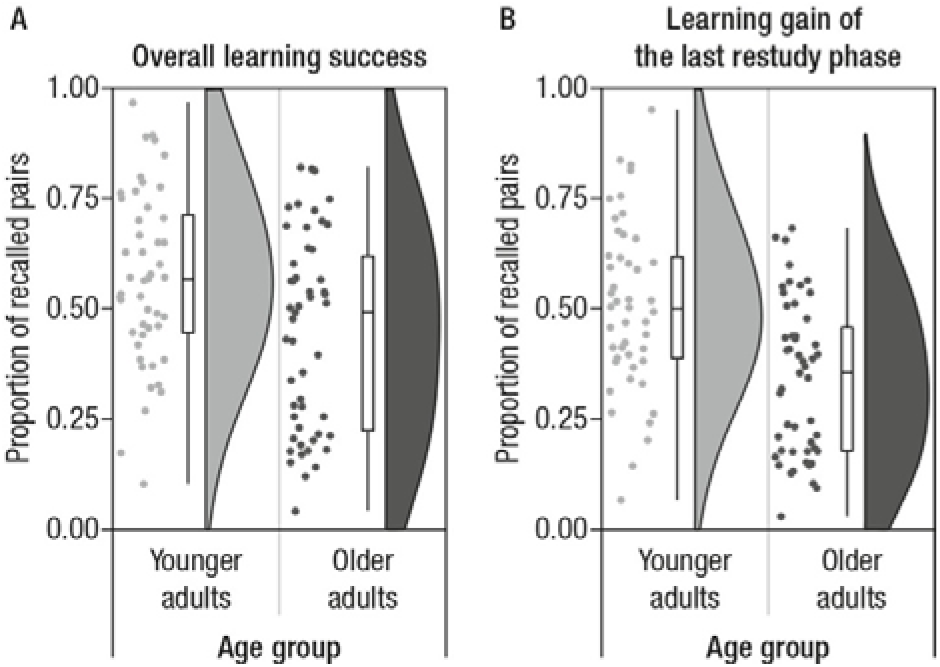
Participants repeatedly studied and recalled scene–word pairs. A. This panel shows their overall learning success as a proportion of recalled pairs at the end of the experiment. Younger adults are shown in light grey and older adults in dark grey. Points represent individual participants, boxplots (median, first, and third quantiles) and violin plots illustrate the sample density. Mean performance levels are close to 0.5 for both age groups with large differences between participants. B. Learning gain of the last restudy phase (i.e., the proportion of recalled pairs in the final recall out of those pairs that had not been successfully remembered in any previous recall phase). Younger adults showed larger gains from restudy than did older adults. This behavioral measure was taken as the basis for the subsequent memory analysis.

### Single-Trial Statistical Analysis

To further investigate the behavioral relevance of modulations in theta and alpha/beta frequencies at the individual level, we extracted single-trial log-transformed power for each participant from the two time-frequency-electrode clusters determined in the first step and averaged across time- and frequency points within the cluster. Single-trial power was then used in a mixed-effects logistic regression (i.e., a generalized linear mixed-effects model, GLMM; Quené and van den Bergh 2008) to predict single-trial accuracy (correct/incorrect, i.e., a binomially distributed response). Alpha/beta power and theta power (both continuous predictors) were z-scored within subjects across trials and centered around the mean of the individual before analysis in order to facilitate the interpretation of parameter estimates. Between-subject differences were included as random effects to account for individual differences in performance level and trial numbers. In order to understand the source of between-person differences in the trial dynamics of alpha and theta power, we included measures of structural integrity for regions of interest, namely cortical thickness of the IFG and HC volume as a between-person fixed effect (continuous predictor, z-scored across the whole sample of younger and older adults). Alpha power modulations have previously been related to the IFG, whereas theta power modulations have been linked to the MTL (Hanslmayr et al. 2011). We therefore allowed IFG cortical thickness to interact with single-trial alpha power and HC volume to interact with single-trial theta power. As we were interested in age differences in SME as well as structure–function relationships, we included age group as a fixed effect and allowed for its interaction (see Figure 2 for an illustration of the mixed-effects model).

Measures of structural integrity enter the analysis as factors explaining the between-person variance. Besides a random subject effect that represents (unknown) differences between individuals, we use the structural measures as fixed effects in our equation to represent some kind of person-specific weighting factor, similar to the effect of age group. Our model thus tests whether there are person-specific factors (namely, the structural integrity of HC and IFG as well as the person’s age group) that modulate the relation between power in a single trial and subsequent memory, i.e., the within-subject effect.

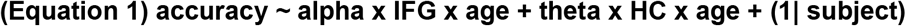

We used maximum likelihood with an Adaptive Gauss-Hermite Quadrature (nAGQ = 10) to estimate model parameters as implemented in the lme4 package (Bates et al. 2015) in R 3.5.2 (R Development Core Team, 2018). We report model parameter estimates with standard errors, z-values, and p-values in Table 1. Note that we run a control analysis including a dummy variable coding for the study in which participants participated (namely the first one, see Fandakova et al. 2018, or the second one, see Muehlroth et al. 2019). This control analysis did not reveal any additional significant effects and is therefore not reported in detail.

**Table 1:**
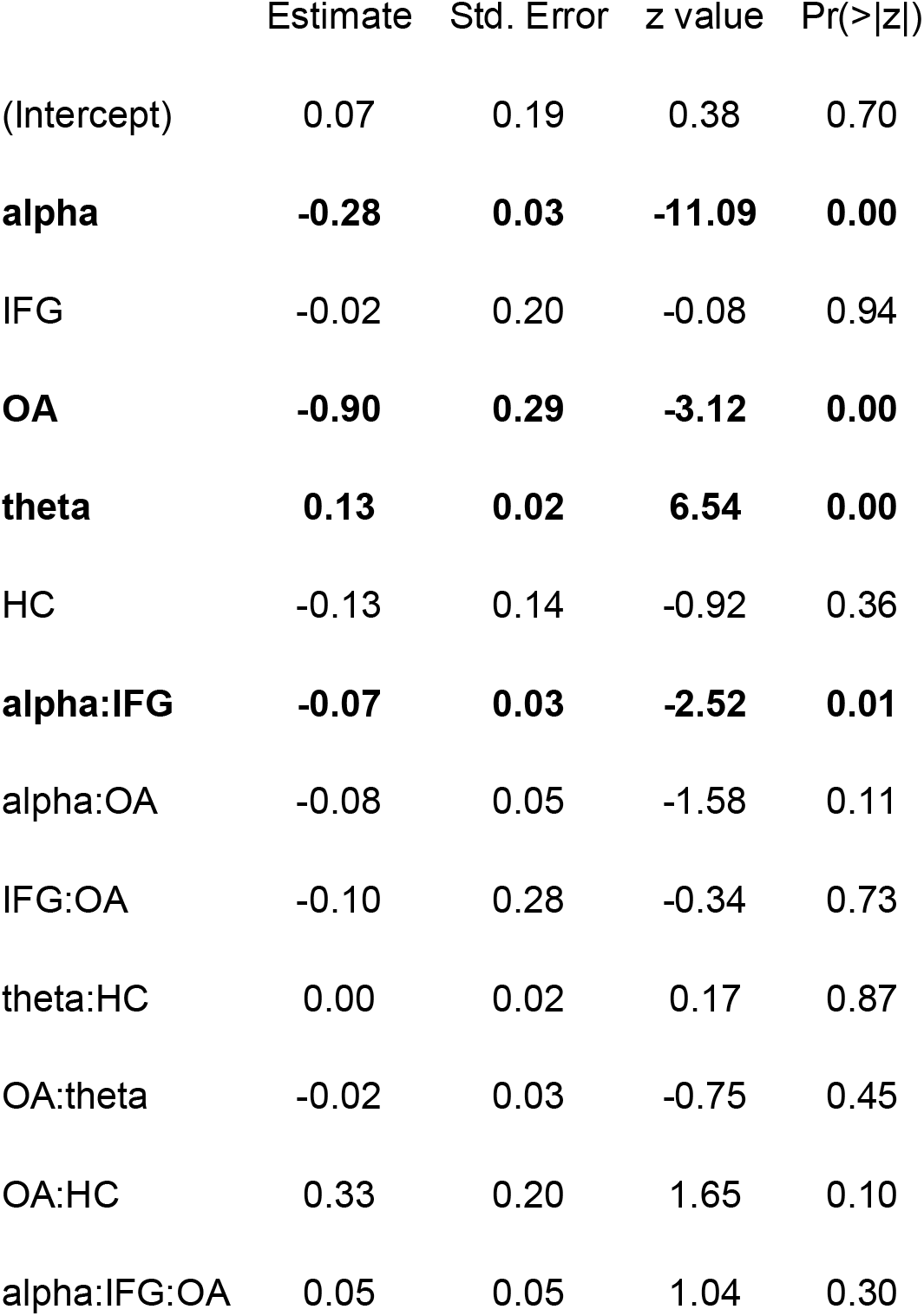
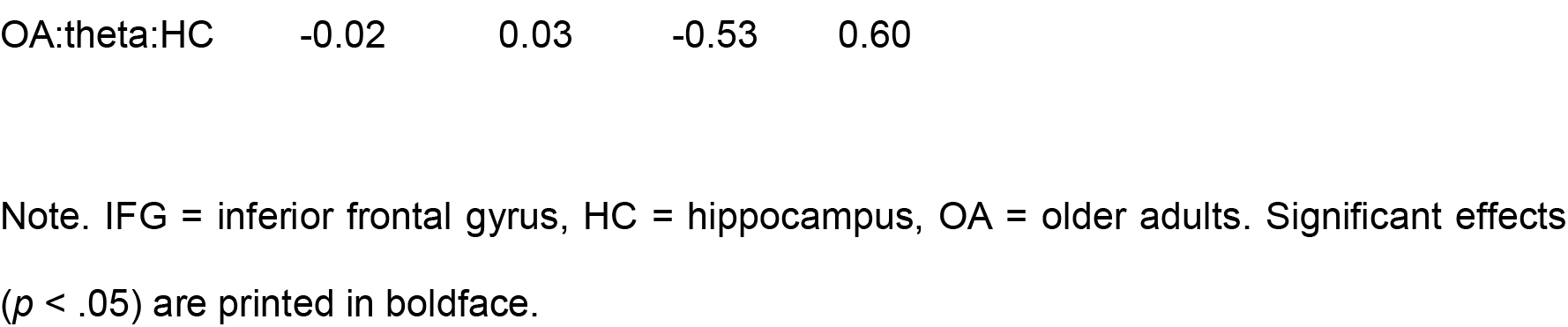
Parameter Estimates for the Mixed-Effects Model Including EEG-SME, Age Group and Measures of Structural Integrity as Predictors of Single-Trial Accuracy

### Code Accessibility

Custom MATLAB and R code of the main analyses are available on https://osf.io/w4pz3/ .

## Results

### Age Differences in Behavior

#### Performance Across the Whole Learning Procedure

Younger and older adults underwent a different learning procedure. Younger adults’ memory was tested twice and older adults’ memory was tested three times. Mean level performances (SD) were: YA_Recall1_ = 0.18 (0.11), YA_Recall2_ = 0.57 (0.20), OA_Recall1_ = 0.03 (0.02), OA_Recall2_ = 0.21 (0.15), and OA_Recall3_ = 0.45 (0.22). We tested for age differences in overall learning success and in the learning gain of the last restudy phase, that is, recall 1 for younger adults and recall 2 for older adults. *Age Differences in Overall Learning Success:* In the final recall test on 440 scene–word pairs for younger and 280 scene–word pairs for older adults, young adults showed higher memory performance than did older adults (*M (SD)* = 0.57 (0.20) vs. *M (SD)* = 0.45 (0.22), *W* = 1590, *p =* 0.010). However, given the large number of study pairs, performance was in a good range and close to the mean level performance of 0.5 in both age groups, thus providing a sufficient number of trials for subsequent memory analyses.

#### Age Differences in Learning Gain in the Last Restudy Phase

In the current study we focussed on the successful learning of scene–word pairs in the last restudy phase. Therefore, we only chose pairs that had not been learned during previous study phases, but were acquired during the last restudy phase (as indicated by successful recall during the final cued recall). We compared them to pairs that were not recalled at any point during the learning procedure. The learning gain in the last restudy phase was higher in younger (*M (SD)* = 0.50 (0.19) than in older adults (*M (SD)* = 0.34 (0.17), *W* = 1782, *p* < 0.001). Thus, while young adults recalled about 50 % of the pairs they did not remember in the previous recall phase, older adults gained less from the restudy phase, despite having an additional opportunity to strengthen their mental image of each pair.

#### Age Differences in Imagery Ratings

To investigate whether older and younger adults differed in their subjective experience as to how well they were able to use the imagery strategy and whether ratings were modulated by subsequent memory, we compared imagery ratings for remembered and not-remembered pairs in younger and older adults (see Figure 4). Both age groups showed a significant effect of subsequent memory on the imagery ratings with higher levels of ratings for subsequently remembered pairs (younger adults: remembered pairs *M (SD)* = 2.38 (.34) vs not-remembered pairs *M (SD)* = 2.05 (.32), *V* = 0, *p* < .001; older adults: remembered pairs *M (SD)* = 2.25 (.52) vs. not-remembered pairs *M (SD)* = 2.06 (.47), *V* = 246, *p* < .001). However, comparing the size of the modulation (computed as the difference in mean ratings for remembered minus not-remembered trials), younger adults showed stronger modulations than did older adults (younger adults: *M (SD)* =.34 (.16) vs. older adults *M (SD)* = .19 (.35), *W* = 1792, *p* < .001).

**Figure 4.**
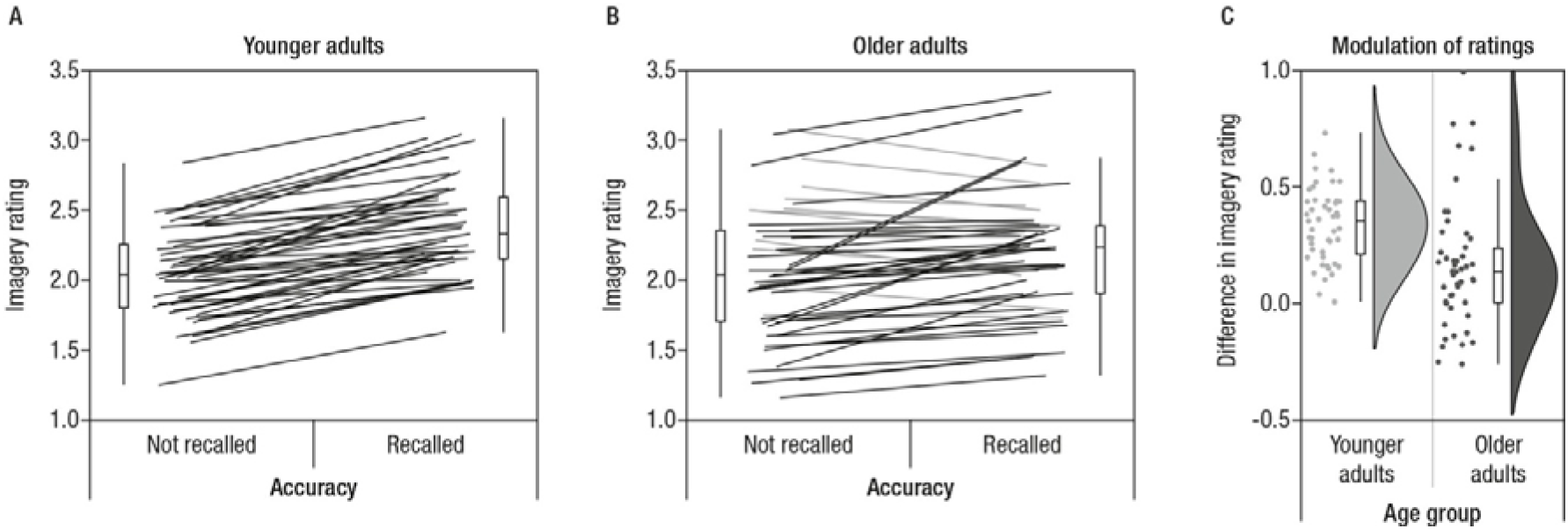
Both younger (A) and older adults’ (B) subjective judgement of the quality of the imagery and elaboration process during encoding varies in accordance to later memory performance (A and B). Paired measures are connected by lines. Participants displaying an effect in the expected direction (i.e., higher ratings for later remembered than for not-remembered pairs) are depicted in black, whereas participants with opposite patterns or no difference are depicted in gray. While the effect was present in all but one younger adult, a larger subsample of older adults did not show the effect. C. Comparing both samples, the modulatory effect was indeed stronger in younger than in older adults.

### Age Differences in Measures of Structural Integrity

Overall, as shown in Figure 5, older adults showed lower (z-normed across the sample) IFG cortical thickness (*M (SD)* = −.92 (0.68)) than did younger adults (*M (SD)* = 0.65 (.66), *t*(97) =11.60, *p* < 0.001). HC body volume (z-normed across the sample) was also reduced in older adults (*M (SD)* = −.41 (0.93)) relative to younger adults, (*M (SD)* = 0.31 (0.96), *t*(97) = 3.80, *p* < 0.001). Despite differences in mean level of structural integrity, variances of the HC and the IFG did not differ between age groups (Fisher’s F-test, *F_HC_*(47,50) = 1.05, *p_HC_* = 0.867 and *F_IFG_*(47,50) = 0.95, *p_IFG_* = 0.867), but variances differed between IFG and HC, with less variance observed for the IFG in both age groups (Fisher’s F-test, F_YA_ (47,47) = 0.48, *p_YA_* = 0.014 and Fisher’s F-test, F_OA_ (50,50) = 0.53, *p_YA_* = 0.028).

**Figure 5.**
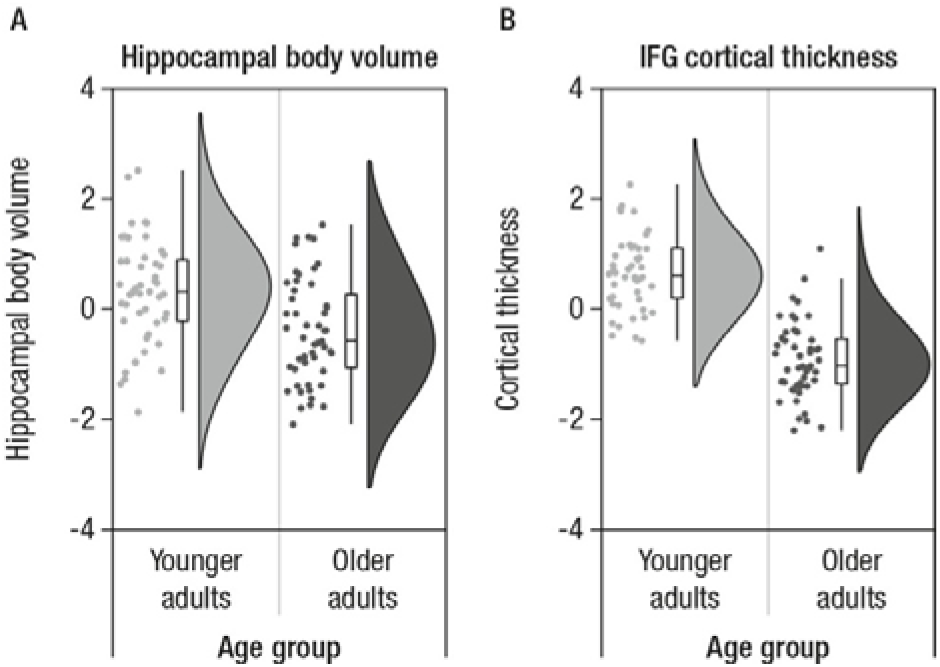
Hippocampal body volume (A) and cortical thickness of inferior frontal gyrus (IFG) (B) shown for each participant (indicated by individual points) together with boxplots and sample density, separated by age group. Older adults (dark grey) show lower IFG cortical thickness and lower hippocampal volume than do younger adults (light grey).

### Age Differences in EEG SME Effects

#### Results of Cluster-permutation Corrected SME Analysis on EEG Data (Averaged on the Subject Level)

Trials with stimuli that were remembered and trials with stimuli that were not remembered during the final cued recall were averaged for each subject. To increase power to detect SME and to derive clusters that are similarly representative for the SME of all participants independent of their age group, grand averages were created by collapsing across age groups. Grand averages were used to determine time-frequency clusters that showed reliable differences between remembered and not-remembered trials on the group level. The cluster-based permutation tests yielded two significant (*p* < .025) clusters of electrodes (see Figure 6): One early cluster (*p* = .024) with a maximum around 500–800 ms that was predominantly found in low frequencies (2–6 Hz), and one later cluster (*p* < .001) with a maximum around 1000–2000 ms encompassing alpha and beta frequencies (8–20 Hz). For ease of comprehension, we refer to the earlier cluster as SME in the theta band and to the later cluster as SME in the alpha/beta frequency band. Both effects displayed a very broad topography. The theta cluster displayed a mid-frontal maximum and the alpha/beta cluster, a centro-posterior maximum.

**Figure 6.**
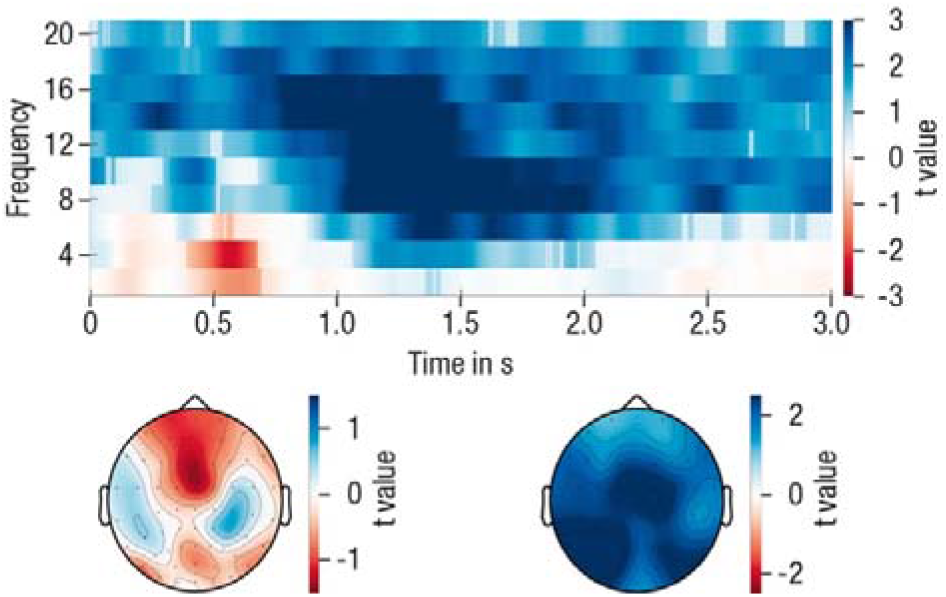
T-values for the comparison of subsequently remembered versus subsequently not-remembered pairs, averaged across electrodes and displayed with their respective topographical distribution. Semi-transparent time-frequency samples are not part of any of the significant clusters. The data were collapsed across participants of both age groups for the derivation of the clusters.

#### Single-Trial Statistical Analysis of EEG SME Effects

To derive a deeper understanding of the theta and alpha/beta power SME, we extracted single-trial log-transformed power for each participant from both time-frequency-electrode clusters determined in the first step. For the purpose of illustration, the SME of the log-transformed and within-person z-transformed theta and alpha/beta power are shown in Figure 7. The effect was present in most participants (indicated by black lines).

**Figure 7.**
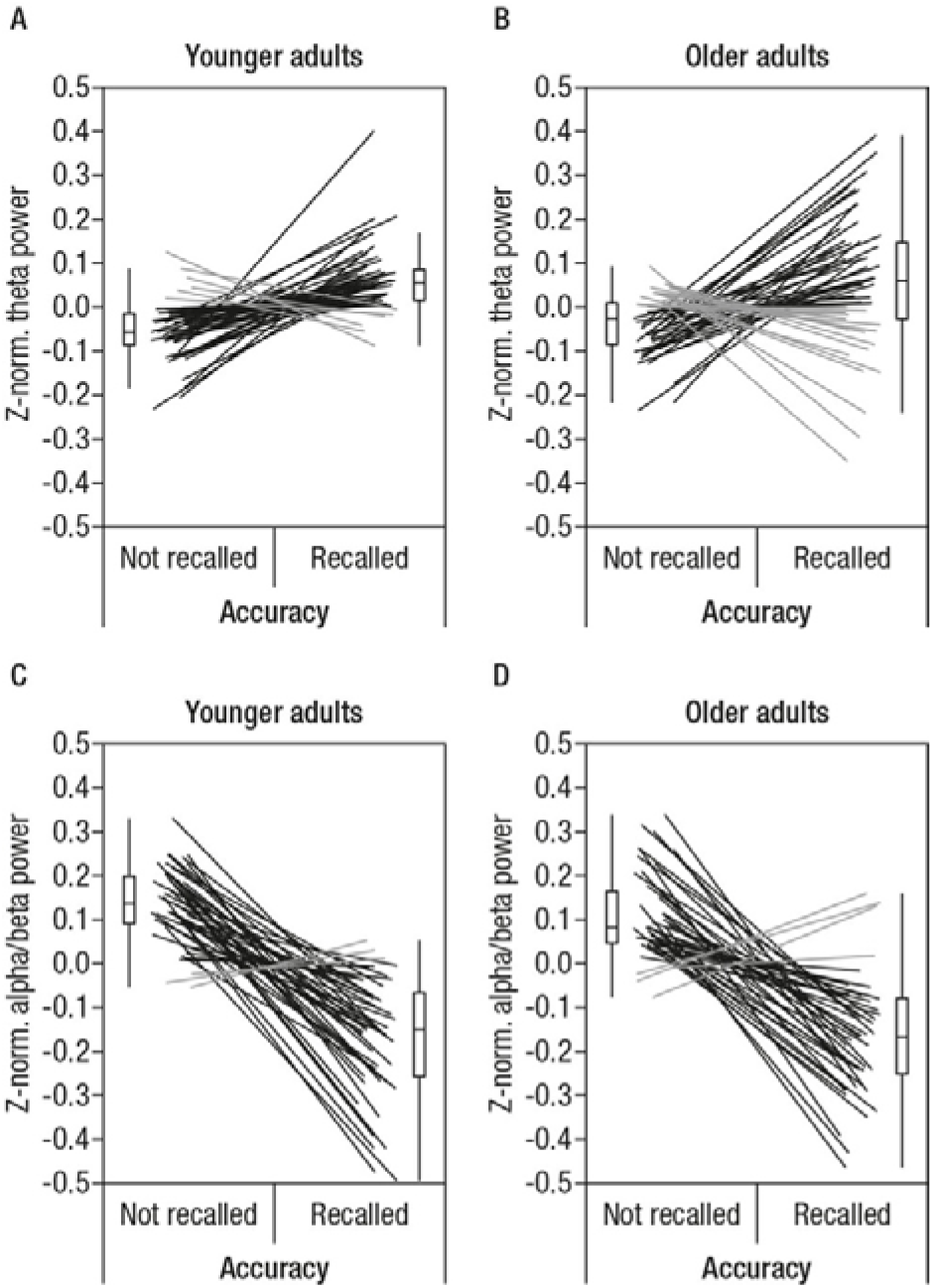
Within-subject modulation of theta power in younger and older adults (A and B, respectively) and of alpha/beta power (C and D) in accordance with subsequent memory. Within-subject z-normalized power was averaged for each participant separately for accurate and inaccurate trials. To display the within-subject effect, data points of individual participants are connected by lines. Participants displaying an effect in the expected direction (i.e, higher power for remembered than not-remembered pairs in theta frequency, and lower power for remembered than not-remembered pairs in alpha/beta frequencies) are depicted in black, whereas participants with opposite patterns or no difference are shown in gray. It is clearly visible that the expected SME were present in most participants in both age groups. Note that this figure is provided only for illustration of the effect formally derived from the cluster-permutation corrected SME analysis on averaged EEG data.

We entered single trial theta and alpha/beta power in a mixed-effects logistic regression to predict single-trial accuracy (correct / incorrect) together with measures of structural integrity of the two brain regions that have previously been related to oscillatory mechanisms of memory formation in the theta and alpha/beta frequency, namely HC and IFG. Thus, we added cortical thickness of IFG and HC body volume as between-person factors and allowed them to interact with the alpha and theta SME, respectively. Finally, we asked whether oscillatory SME and structure-function relationships would differ between age groups. We therefore included age group as an additional predictor in our model, allowing for interactions with all other predictors. The model had a conditional R^2^ = 0.25, thus, our predictors accounted for 25 % of the variance in single-trial accuracy. All parameter estimates can be found in Table 1.

Single trial alpha/beta and theta power were robustly linked to performance (both *p* < .001), with higher theta power and lower alpha/beta power yielding a higher likelihood for correct recall. Importantly, although age group as a fixed effect was a strong predictor of performance (*p* = .002), consistently with the behavioral results reported above, age did not interact with any of the other predictors. These results suggest that SME in alpha/beta and theta power were similar across age groups. To further illustrate the similarity between younger and older adults with regard to these within-subjects power modulations, we displayed the predicted probabilities separately for younger and older adults in Figure 8.

**Figure 8.**
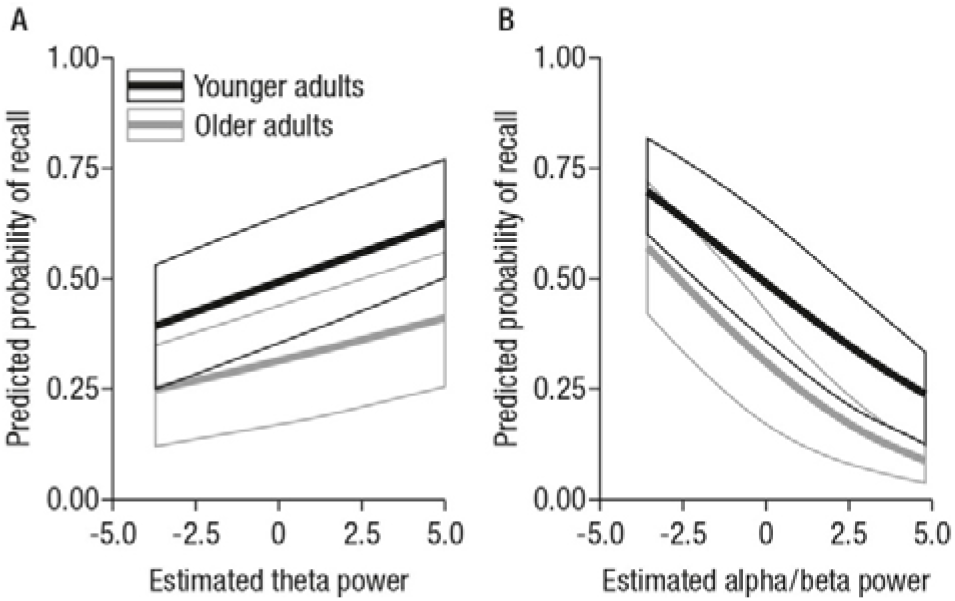
Main effects of theta (A) and alpha/beta (B) power on the predicted probability of recall. While the mixed-effects model showed a main effect of age group, there were no interactions with alpha/beta or theta power. Nevertheless, we display the predicted probabilities separated by age group to illustrate this point.

HC volume neither predicted accuracy (*p* = .36) nor showed significant interactions with theta power (*p* = .87). However, the effect of alpha power on the probability of successful recall was modulated by IFG cortical thickness (*p* = .02). Accordingly, for participants with lower cortical thickness, modulations in alpha power less reliably predicted subsequent memory performance (see Figure 9). Importantly, this structure–function relationship did not differ by age group (*p* = .30), underscoring that basic mechanisms of memory formation as well as the factors underlying interindividual differences in memory performance were similar in younger and older adults. Nevertheless, given the above reported age differences in structural integrity of the IFG, most participants of the lower quantiles happen to be older adults and thus have a higher probability for a less reliable relation between alpha power modulations and subsequent memory performance (as displayed in Figure 9).

**Figure 9.**
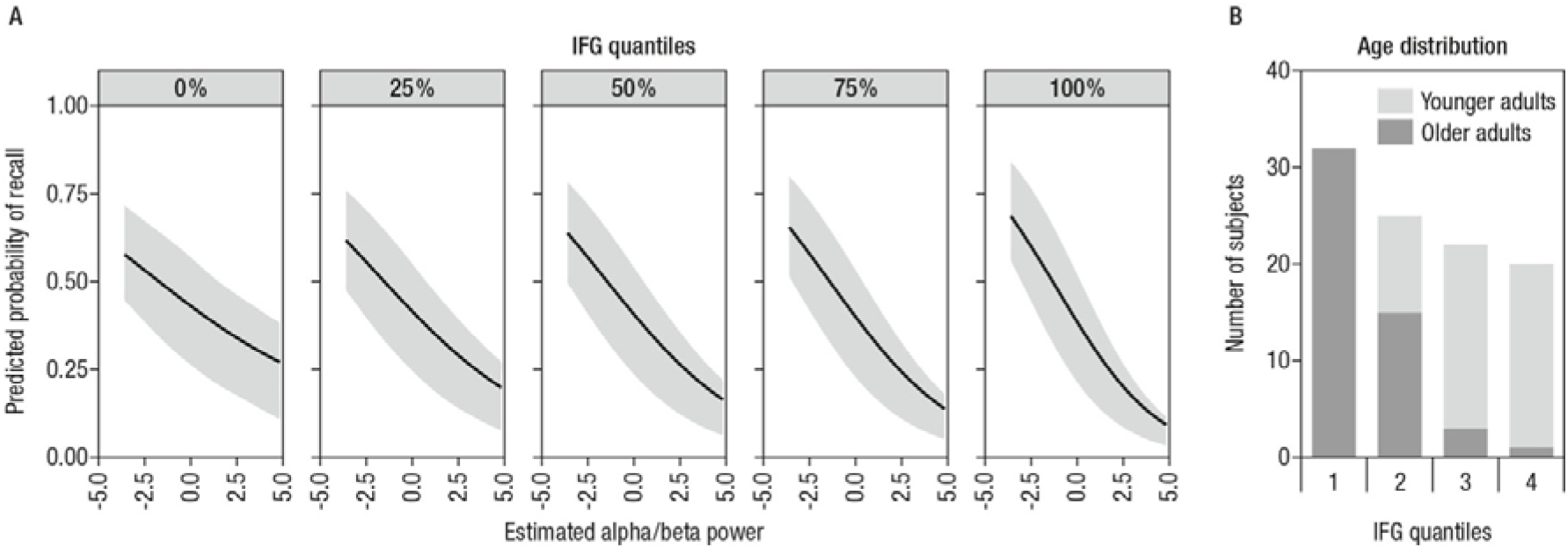
A. The effect of alpha power on predicted probability of successful recall is modulated by IFG cortical thickness, as shown by displaying predicted probabilities of varying alpha power for different IFG quantiles. For participants with lower cortical thickness, modulations in alpha power less reliably predicted subsequent memory performance. B. Distribution of older and younger adults across different levels of structural integrity of the inferior frontal gyrus (IFG) (represented by quantiles).

## Discussion

We set out to investigate SME in oscillatory activity in young and older adults and their relation to age differences in structural integrity of key brain regions for memory formation. We found that single-trial alpha/beta and theta power were reliable predictors of memory success or failure in a cued-recall task in both younger and older adults. The observation of similar <u>within-person</u> power modulations (i.e., SME) indicate that basic mechanisms of memory formation do not differ between age groups, thus, low alpha/beta power and high theta during encoding increases recall probability in both age groups. We then examined whether differences in the structural integrity of the IFG, a brain region closely linked to elaborative processes during encoding, and the HC, a brain region relevant for the binding of information into a coherent memory representation, could explain <u>between-person</u> differences in oscillatory mechanisms of memory formation in the alpha/beta and theta band, respectively. Thus, we asked whether a person with higher structural integrity (compared to others) would also show stronger functional modulations of oscillatory activity. We indeed found that cortical thickness of the IFG was related to SME in the alpha/beta band. For participants with greater cortical thickness of the IFG, a difference in alpha power was a better predictor of subsequent memory performance than for participants with lower cortical thickness. In contrast to our hypothesis, we did not observe effects of HC volume on oscillatory dynamics in the theta band. Importantly, while we observed overall age differences in memory accuracy as well as in the structural integrity of the IFG and the HC, we did not find an age-differential effect for the observed structure–function relationship between alpha power and cortical thickness of the IFG. However, older adults were more frequently represented among the participants with low cortical thickness and consequently weaker SME in the alpha band. Thus, our results suggest that differences in the structural integrity of the IFG are the basis not only for interindividual differences, but also for age differences in memory formation.

### Contributions of structural integrity to oscillatory mechanisms of successful memory formation

Episodic memory formation is tightly linked to interactions between MTL regions that bind incoming information into coherent representations and PFC regions that select and elaborate these representations (Simons and Spiers 2003; Shing et al. 2010). These two systems often show opposing oscillatory behavior – desynchronization in the alpha band supports successful memory elaboration (Hanslmayr et al. 2016), whereas synchronization in the theta band mediates binding (Staudigl and Hanslmayr 2013). With regard to the latter, note that some studies also reported that decreases in theta power were predictive for subsequent memory performance (for an overview, see Hanslmayr and Staudigl 2014).

In our study, we observed reliable SME in the theta band with increased power for scene–word pairs that were later successfully remembered compared to those that were not remembered. Our finding is in line with studies using intracranial recordings that found theta power increases during successful encoding in the HC (Lin et al. 2017; for similar results see also Sederberg et al. 2003; Lega et al. 2012). Similarly, SME in the theta band were previously observed in young adults with their source being located to the MTL and ventral lateral regions (Hanslmayr et al. 2011). These findings support the idea that the MTL, and in particular the HC, plays a critical role for episodic memory formation via the integration of multiple features into coherent memory traces. This assumption was further underlined by a recent magnetoencephalography (MEG) study (Staudigl and Hanslmayr 2013) that found theta–gamma coupling in the MTL during item–context binding in episodic memory. Based on these findings, we therefore hypothesized that interindividual differences in the structural integrity of the HC modulate SME in the theta band. However, we did not find strong evidence for this assumption. Since our conclusions are based on EEG scalp recordings, it is possible that the observed effects in the theta frequency do not directly capture HC activity, but rather reflect HC–frontal interactions during encoding (for review, see Klimesch 1999; Nyhus and Curran 2010). Of note, despite a localization of the theta band SME to the broader MTL region, the study by Hanslmayr et al. (2011) also did not find reliable single-trial correlations between theta power and the BOLD signal in the region. Thus, neither functional activation of the MTL (as in Hanslmayr et al. 2011) nor HC volume alone seems to be a good predictor for (M/EEG-) theta power modulations.

In contrast to the observed *increases* in power in the theta frequency range, we observed reliable alpha/beta power *reductions* for scene–word pairs that were successfully remembered as compared to those that were not remembered. Reduced alpha power for items later remembered has previously been found in EEG studies using young adult samples (Fellner et al. 2013; Noh et al. 2014). The observation of reduced alpha power for successful memory formation is in line with recent theoretical accounts (Hanslmayr et al. 2016; Hanslmayr et al. 2012) suggesting that information processing capacity can be increased within local cell assemblies via a decrease in local synchronization. Thus, long time windows of desynchronization in the alpha/beta frequency range may indicate prolonged elaborative encoding that in turn facilitates episodic memory success. In our study, participants were instructed to use an imagery strategy during encoding. In line with electrophysiological SME, imagery ratings also differed according to subsequent accuracy: Recalled pairs received higher imagery ratings in both age groups, underlining that successful deep elaboration during encoding helps memory formation (Craik and Lockhart 1972; Craik and Rose 2012). This assumption is further supported by our finding that the structural integrity of the IFG modulated the contribution of alpha power to successful memory performance. Many studies have implicated the IFG in successful memory formation, particularly for semantic elaboration (for reviews see Kim et al. 2011; Paller and Wagner 2002). Our results resonate with a previous EEG-MRI study demonstrating that beta power decreases correlated with increases in the BOLD signal in the left IFG on a trial-by-trial basis (Hanslmayr et al. 2011). The importance of the IFG for oscillatory desynchronization was further demonstrated by a recent study that used TMS to interfere with memory formation (Hanslmayr et al. 2014): The authors were able to demonstrate a behavioral impairment that was selective for beta frequency stimulation of the left IFG. Our study nicely completes this picture by demonstrating that *structural* integrity of this region is relevant for the modulatory effects of alpha/beta power on successful memory formation, revealing a clear structure–function relationship that predicts behavior. The use of an intentional memory paradigm in the current study probably strengthened the observation of the relation between structural integrity and the modulatory effect of alpha power, since the instruction fostered a deep, semantic elaboration which is known to recruit the IFG in particular.

### No age differences in neural mechanisms of memory formation?

Importantly, we found that in older adults, similarly to younger adults, successful memory encoding was accompanied by reliable modulations of theta and alpha/beta power. Put differently, SME, which are per definition <u>within-person</u> effects, were not modulated by age, despite overall age differences in performance, indicating that the basic mechanisms of memory formation do not differ between age groups. In addition, the observed structure– function relationship between IFG and alpha power with its effect on memory performance did not differ by age group, despite overall age differences in cortical thickness of the IFG.

Our results are in line with previous fMRI studies (de Chastelaine et al. 2011, 2016; Shing et al. 2016) that observed robust SME in the IFG and the HC in both younger and older adults. Similarly, a meta-analysis of the subsequent memory paradigm in age-comparative settings (Maillet and Rajah 2014) came to the conclusion that MTL and IFG are among those brain regions that show age-invariant patterns of SME, at least with regard to fMRI. Our study shows a parallel age invariance in oscillatory SME, thereby adding another piece of evidence to what has been established using fMRI.

To the best of our knowledge, there is only one recent study that investigated oscillatory SME in younger and older adults (Strunk and Duarte 2019). Similar to our results, they found SME in alpha/beta and theta bands that did not differ by age group. Importantly, in contrast to their study, which used recognition memory, we used cued recall to test successful memory formation. This procedure has clear advantages over recognition tasks in which hits that were committed with high confidence are frequently contrasted with low-confidence hits collapsed together with misses (Strunk and Duarte 2019). Whereas recalling an associate in a cued-recall task is a clear indication of recollection, hits in recognition tasks do not only rely on recollection, but also on familiarity, which is only partly taken into account by counting low-confidence hits as not-remembered. Furthermore, the reliance on confidence ratings can be particularly problematic in age-comparative settings, as metacognitive differences between younger and older adults may affect the subsequent memory analysis (e.g., older adults appear to commit false alarms with high confidence, see (Shing et al. 2009; Fandakova et al. 2013). In addition, in our study, participants were instructed in, and practiced using an imagery strategy that fosters associative and elaborative processing. Our choice of an intentional encoding task was motivated by the observation that cognitive strategies that are spontaneously adopted often differ between age groups (Rugg and Morcom 2005). However, differences in encoding strategies (e.g., shallow versus deep encoding) have been shown to modulate SME (Hanslmayr and Staudigl 2014). The explicit strategy instruction in our study therefore aimed to minimize the possible confound of neural differences with age group differences in cognitive strategies. Indeed, like younger adults, older adults’ subjective judgement of elaboration success varied with subsequent memory performance. By using an intentional encoding task that fostered elaborative processing, we may have successfully induced effective encoding strategies to improve episodic memory performance in older adults, which then manifest as age-invariant mechanisms of memory formation.

At the same time, as expected, older adults’ memory performance was overall significantly lower than younger adults’ performance. How can age-invariant mechanisms of memory formation (i.e., similar SME in younger and older adults) be reconciled with the well-known general age differences in memory performance? First, it is important to note that SME are per definition <u>within-person</u> differences, thus, driven by a neural mechanism that predict whether encoding will be successful or not – in a given person. Age differences in memory performance, on the other hand, are per definition between-person differences referring to a stable rank order between persons of different ages. Between-person differences and within-person mechanisms can be identical, but do not necessarily have to be identical (for a theoretical treatment of this issue, see for example Gayles and Molenaar 2013; Nesselroade et al. 2007; Voelkle et al. 2014). The current study focused on age differences in within-person mechanisms, thus, SME in younger and older adults, and examined whether these would be modulated by age differences in structural integrity.

First, it is notable that there were reliable age differences in structural integrity in IFG and HC, in line with previous reports on differences in volume and cortical thickness in these brain regions (Raz and Rodrigue 2006). Age-related structural changes in these regions have previously been linked to episodic memory performance and are regarded as underlying functional activation differences between younger and older adults (for a review, see Nyberg 2017). However, while it is reasonable to assume that structural integrity of a brain region modulates its functional activation and thereby influences behavioral outcomes, this triadic relation has not been sufficiently established. In the current study, we observed that structural integrity of the IFG modulated SME in the alpha/beta. Thus, our results support the view that functional activation and structural integrity are closely related by revealing the contribution of structural integrity of the IFG to oscillatory power modulations related to memory success. While the observed structure–function relationship was independent of age, importantly, the participants with low cortical thickness were mostly older adults and consequently, power modulations in the alpha/beta band were also less predictive for subsequent memory performance in these older adults. Thus, our findings suggest that the reduced behavioral performance in older adults can be attributed to lower IFG thickness, which is associated with smaller alpha SME. An altered slope of the alpha power function is in line with the prominent hypothesis of an overall noisier system in older adults (Li et al. 2000) that has far-reaching consequences for performance (Garrett et al. 2013).

Notably, brain regions beyond IFG and HC also contribute to successful episodic memory encoding and are also affected by senescent-related changes (Ankudowich et al. 2016, 2019; Fjell et al. 2016). Interindividual differences in the onset and severity of senescent changes may already lead to differences in episodic memory performance in middle-aged adults (Park et al. 2013). Furthermore, it is reasonable to assume that the neural integrity of key memory regions may also modulate activity in distally distinct parts of the network (see Maillet and Rajah, 2013, for a review, and recent evidence by Becker et al. 2019 on covariance patterns between MTL volume and functional activation in IFG). Future studies are required to better understand brain-wide structure–function relations underlying successful memory performance.

To conclude, our results support the assumption that oscillatory mechanisms of successful memory formation do not differ between younger and older adults. At the same time, they suggest that age-related differences in structural brain integrity contribute to the decline of episodic memory performance in older adults, since we observed that structural integrity of the IFG modulates oscillatory SME in the alpha/beta band. Thus, our findings suggest that the reduced behavioral performance in older adults can be attributed to lower IFG thickness, which is associated with smaller alpha SME.

While it has to be kept in mind that our sample of older adults are not necessarily representative for their peers, but represent high-performing, healthy seniors, our findings further support the continuously mounting evidence that the maintenance of structural integrity goes hand in hand with the maintenance of youth-like mechanisms of memory formation in older adults (Nyberg et al. 2012; Fandakova et al. 2015; Nyberg and Pudas, 2019).

## Acknowledgments

We thank all our student assistants for their support in data collection. We thank Julian Kosciessa for helping with programming the task, Attila Keresztes for help with processing of the structural MRI data and Beate Mühlroth and Xenia Grande for their support in data collection. We are also grateful to Julia Delius for editorial assistance.

